# Spatial Dynamic Subspaces Encode Sex-Specific Schizophrenia Disruptions in Transient Network Overlap and its Links to Genetic Risk

**DOI:** 10.1101/2023.07.18.548880

**Authors:** A. Iraji, J. Chen, N. Lewis, A. Faghiri, Z. Fu, O. Agcaoglu, P. Kochunov, B. M. Adhikari, D.H. Mathalon, G.D. Pearlson, F. Macciardi, A. Preda, T.G.M. van Erp, J. R. Bustillo, C. M. Díaz-Caneja, P. Andrés-Camazón, M. Dhamala, T. Adali, V.D. Calhoun

## Abstract

**Background:** Recent advances in resting-state fMRI allow us to study spatial dynamics, the phenomenon of brain networks spatially evolving over time. However, most dynamic studies still use subject-specific, spatially-static nodes. As recent studies have demonstrated, incorporating time-resolved spatial properties is crucial for precise functional connectivity estimation and gaining unique insights into brain function. Nevertheless, estimating time-resolved networks poses challenges due to the low signal-to-noise ratio, limited information in short time segments, and uncertain identification of corresponding networks within and between subjects.

**Methods:** We adapt a reference-informed network estimation technique to capture time-resolved spatial networks and their dynamic spatial integration and segregation. We focus on time-resolved spatial functional network connectivity (spFNC), an estimate of network spatial coupling, to study sex-specific alterations in schizophrenia and their links to multi-factorial genomic data.

**Results:** Our findings are consistent with the dysconnectivity and neurodevelopment hypotheses and align with the cerebello-thalamo-cortical, triple-network, and frontoparietal dysconnectivity models, helping to unify them. The potential unification offers a new understanding of the underlying mechanisms. Notably, the posterior default mode/salience spFNC exhibits sex-specific schizophrenia alteration during the state with the highest global network integration and correlates with genetic risk for schizophrenia. This dysfunction is also reflected in high-dimensional (voxel-level) space in regions with weak functional connectivity to corresponding networks.

**Conclusions:** Our method can effectively capture spatially dynamic networks, detect nuanced SZ effects, and reveal the intricate relationship of dynamic information to genomic data. The results also underscore the potential of dynamic spatial dependence and weak connectivity in the clinical landscape.

## Spatially Dynamic Analyses in rsfMRI: Quantifying Spatial Network Coupling

The human brain maintains, regulates, adapts, and responds to a rich repertoire of behavior and mental activities via the continuous reconfiguration of coordinated intrinsic activities. At a large scale, these coordinated intrinsic activities are thought to manifest as a set of discrete yet interactive neuronal assemblies, commonly referred to as functional units or functional sources [1].

This view has gained traction in the field of resting-state functional magnetic resonance imaging (rsfMRI), where spatially fixed nodes or data-driven estimations of functional sources, e.g., functional networks [2-4] or functional parcels [5-7], have been used to model the functional interactions among functional units.

However, rsfMRI studies are often limited by the assumption that spatial patterns of functional sources remain constant over the length of the scan. This assumption has influenced the research, as most functional connectome and chronnectome [8] studies use average voxel time series of fixed spatial regions to estimate the time courses of functional sources and compute whole-brain static or temporally dynamic functional connectivity (FC).

It is important to consider both temporal and spatial dynamics. Temporal dynamics refer to variations in the temporal patterns of functional units, commonly assessed through changes in second-order statistics. Spatial dynamics, on the other hand, refer to variations in the spatial distribution of a source over time [1, 9-12]. The continuous reconfiguration of coordinated intrinsic activities can result in changes in the spatial patterns of functional units over time. Therefore, relying solely on the average time series over anatomically fixed regions, which overlooks the temporal evolution of functional units, leads to suboptimal FC estimation and imprecise inferences.

Spatial dynamics also carry unique information about brain function hidden from existing spatially-static approaches. This includes the spatial coupling and uncoupling of functional units over time. Previous work [9] has shown that brain networks can dynamically segregate and integrate in space, including the transient emergence of the cerebellar and primary visual networks within the spatial patterns of other brain networks. Nevertheless, rsfMRI studies have often treated functional units as static entities that are non-overlapping or spatially independent. Consequently, the crucial aspect of their spatial dependence and its temporal changes have often been overlooked.

Spatial dependence is a crucial aspect of statistical analyses of spatially distributed data. However, its significance in the investigation of brain function has not received sufficient attention, despite its essential role in understanding the merging and dissociation of functional units over time. This dynamic spatial dependence reflects the interplay between integrative and specialized processes, and it can be used to explore the balance between integration and segregation in brain networks.

Here, we use spatial functional network connectivity (spFNC) to describe the spatial dependence between networks, consistent with the definition of temporal functional network connectivity, which refers to the temporal dependence between networks. By using spFNC, we can gain insights into the underlying mechanisms of brain function and its adaptability across different states.

### Capturing Time-Resolved Network-Specific Spatial Patterns

To effectively estimate time-resolved spFNC (tr-spFNC), it is essential to first obtain precise spatial patterns of networks in a time-resolved manner. We introduce a time-resolved reference-informed network estimation approach that derives time-varying spatial maps for each network while controlling for the impact of other networks (**Fig. 1**). This approach also overcomes the uncertainty associated with post hoc matching procedures, which can be more problematic in a time-resolved setting [4, 13]. Additionally, by controlling for other networks, we can disentangle the specific contributions of each network and accurately capture their spatial patterns over time. Furthermore, the combination of reference-informed and spatial-constraint mechanisms can effectively address the challenges posed by the low signal-to-noise ratio (SNR) and limited information in small time segments of rsfMRI. Spatial constraints, informed by prior reference knowledge about the spatial distribution of functional units, restrict the search space and act as regularizers. This mitigates overfitting to noise, allowing for the capture of the underlying signal and ultimately improving the reliability of network estimations within a shorter time frame.

**Fig. 1.**
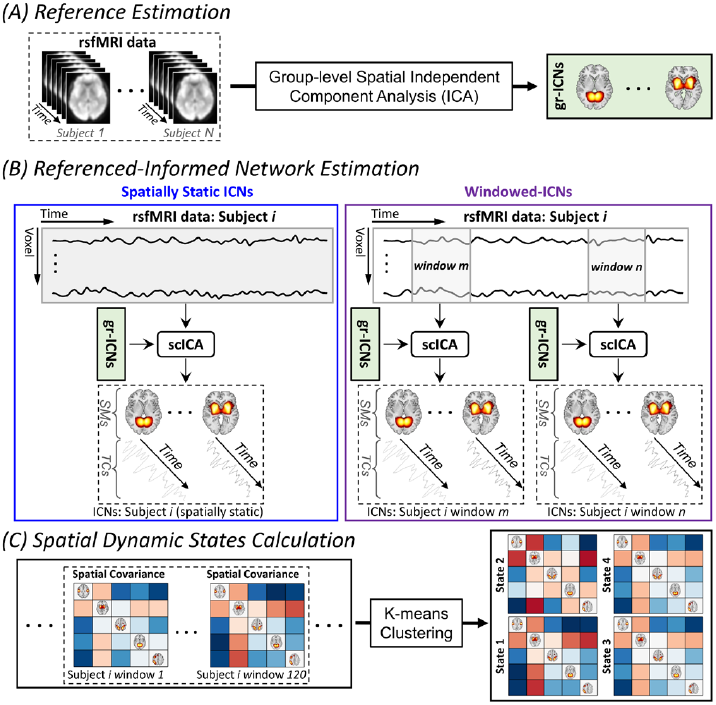
Schematic of the Analysis Pipeline. (A) Using group-level spatial independent component analysis (ICA) [14] to obtain group-level intrinsic connectivity networks (gr-ICNs) as a functional unit reference. (B) Applying spatially constrained ICA (scICA) to estimate the correspondence of gr-ICNs to estimates from a given subject. The left panel shows the standard scICA application to estimate spatially-static ICNs (i.e., assume spatial patterns of ICNs remained fixed over time), and the right panel shows the proposed approach to estimate time-resolved ICNs information as windowed-ICNs. (C) Calculating whole-brain spatial dynamic states from spatial covariance matrices.

We first performed group-level spatial independent component analysis (gr-spICA) [14] with a model order of 20 [9, 15] using the Group ICA of FMRI Toolbox (http://trendscenter.org/software/gift) [16] and obtained large-scale brain networks used as the templates for downstream analysis (**Fig. 1(A)**). Fourteen components with very high ICASSO stability index (average ± standard deviation = 0.96 ± 0.01, minimum-maximum = 0.93-0.98) were identified as brain networks based on their temporal and spatial properties and knowledge from previous studies [4, 9, 15]. They include the primary and secondary visual (VIS-P/VIS-S), primary and secondary somatomotor (MTR-P/MTR-S), subcortical (SUB), cerebellar (CER), attention (ATN), frontal (FRNT), left and right frontoparietal (FPN-L/FPN-R), posterior and anterior default mode (DMN-P/DMN-A), salience (SN), and temporal (TEMP) networks (**Fig. 2(A)**).

**Fig. 2.**
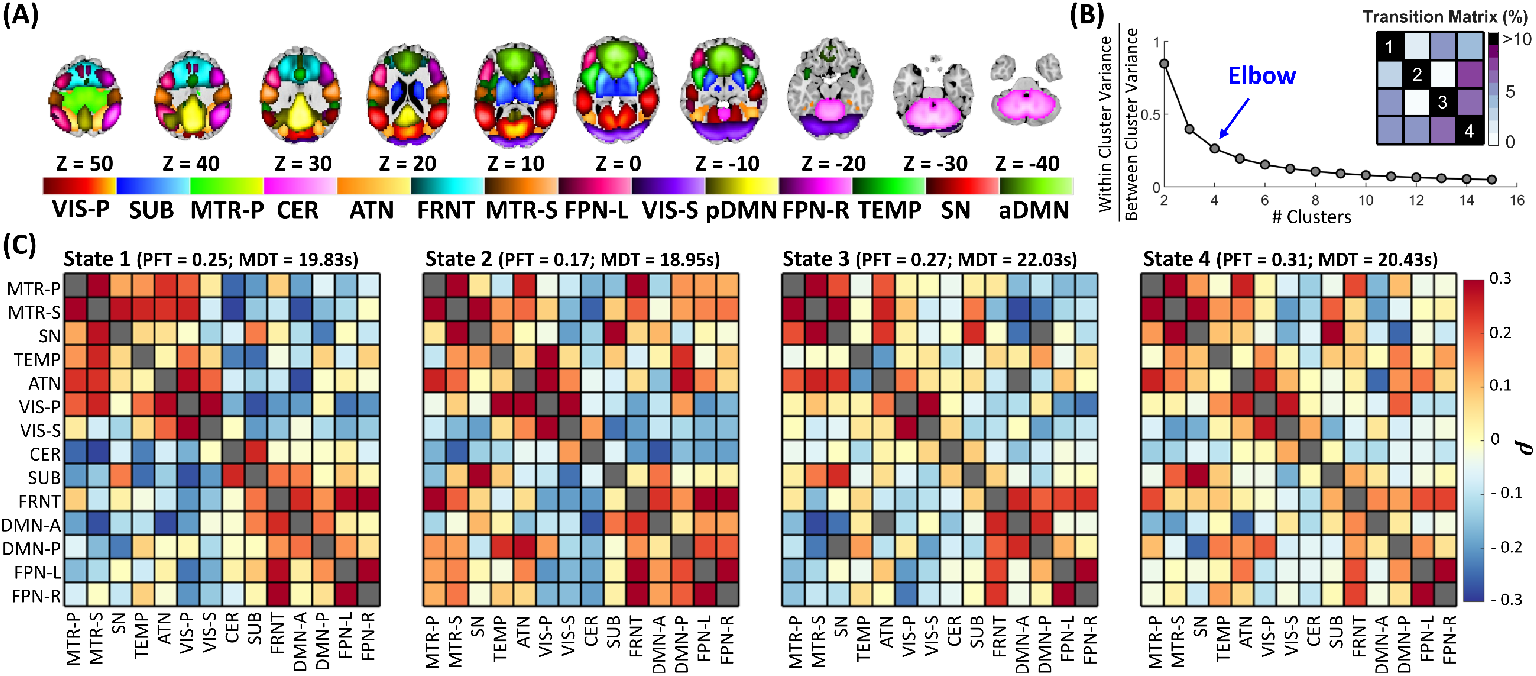
Capturing Global Brain States Dynamics using Low-dimensional Spatial Dynamic States. (A) Visualization of the intrinsic connectivity networks (ICNs). Each color in the composite map represents the spatial map of one ICN thresholded at |Z| > 1.96 (p = 0.05). (B) Estimation of the optimal number of states. The *k*-means clustering procedure was conducted for cluster numbers from 1 to 15 clusters. The ratio of within-to between-cluster variance was calculated for each clustering, and the elbow criterion was used to estimate the number of global states. (C) The four spatial dynamic states are identified using *k*-means clustering with L1 distance. The fraction rate (FT) is the fraction of times a subject spends in a given state, and mean dwell time (MDT) represents the average time a given subject stays in a given state before switching to another state. The MDT is similar across states (18.95 ∼ 22.03 seconds(s)), while FT shows more difference across states (0.17 ∼ 0.31).

Next, we combined a spatially constrained ICA (scICA) method called multivariate-objective optimization ICA with reference (MOO-ICAR) [17] (**Fig. 1(B)**: **Left**) and the sliding-window technique [16] to estimate time-resolved networks corresponding to the templates (**Fig. 1(B)**: **Right**). MOO-ICAR has been shown to perform well in capturing sample-specific information for different data lengths and intrinsic connectivity networks (ICNs) [4] and is also robust to artifacts [18].

The sliding-window technique is the most commonly used technique to study brain dynamics due to its simplicity, ease of use, and substantial similarity with the conventional FC procedure, making the interpretation of findings straightforward [16]. The sliding-window technique was implemented using a tapered window created by convolving a rectangle (width = 60 seconds) with a Gaussian (σ = 6 seconds) and a sliding step size of one [16, 19]. Prior to estimating brain networks, we cleaned each voxel time course using detrending, despiking, removal of motion effects, and filtering to reduce the impact of noise and nuisance signals [9].

### Low-dimensional Spatial Dynamic States Encapsulates Global Brain State Dynamics

We quantified tr-spFNC by calculating the spatial covariance of the large-scale networks at each time window. An increase and decrease in spatial covariance indicate integration and segregation between networks. We next captured the global brain state dynamics by estimating a set of reoccurring distinct spFNC patterns that represent dynamic states [16]. Following previous work and recommendations [19, 20], we applied *k*-means clustering using L1 distance and elbow criterion (**Fig. 2(B)**) and identified four spatial dynamic states (**Fig. 2(C)**).

The fraction of time that individuals spend in spatial dynamics states varies significantly, with State 4 having an approximately twofold higher fraction rate (FT) compared to State 2 (0.31 vs. 0.17). State 4 demonstrates the lowest level of overall network integration, while States 1 and 2 show the highest integration. Conversely, the mean dwell time (MDT), which indicates the amount of time spent at each state per visit, is very similar across all states ranging between 18.95 to 22.03 seconds. In other words, while the life expectancy of spatial dynamic states (i.e., MDT) is similar on average, the total amount of time the brain stays in each state varies.

### The Clinical Relevance of Dynamic Spatial Coupling: A Schizophrenia Study

Schizophrenia has been hypothesized as a disconnection syndrome, where functional integration disruptions are thought to have a more profound impact on behavior and psychopathology than aberrations in single brain networks [21, 22]. Therefore, identifying these disruptions is crucial to decipher the underlying neurobiology of schizophrenia. In this study, we employed tr-spFNC and spatial dynamic states to investigate alterations in the continuous adaptive reconfiguration of functional integration and segregation associated with schizophrenia.

We hypothesized that there would be schizophrenia-related changes in the dynamic spFNC of large-scale networks, including the SN, pDMN, FPNs, ATN, TEMP, MTRs, SUB, and CER, based on our prior spatial dynamic analysis [9], which revealed schizophrenia-related alterations in their spatial maps. Alterations in the functional connectivity of these networks have been frequently reported in previous work [10, 20, 23-30]. We posited sex-specific disruption in spFNC between the pDMN and SN, given their abnormal functional connectivity has been linked to negative symptoms, which have been shown to be more pronounced in men with schizophrenia compared to women [25, 31-35]. Furthermore, this functional connectivity mediated the association between sex and mental rotation [36], potentially explaining the frequently observed schizophrenia-by-sex interactions in the mental rotation task [35]. Additionally, previous research has shown sex differences in the functional connectivity of these networks in youths with Autism spectrum disorder [37], which shares significant clinical and genetic components with schizophrenia [38].

For this purpose, we used 3 Tesla rsfMRI data from 508 subjects (Table 1) and conducted statistical comparisons using a generalized linear regression model (GLM). We included age, sex, mean framewise displacement (meanFD), and site as confounding factors and diagnosis and sex-by-diagnosis interactions as predictors of interest. We ran a GLM separately for each spFNC pair of each spatial dynamic state. The p-values are corrected using the 5% false discovery rate (FDR) [39].

**Table 1.**
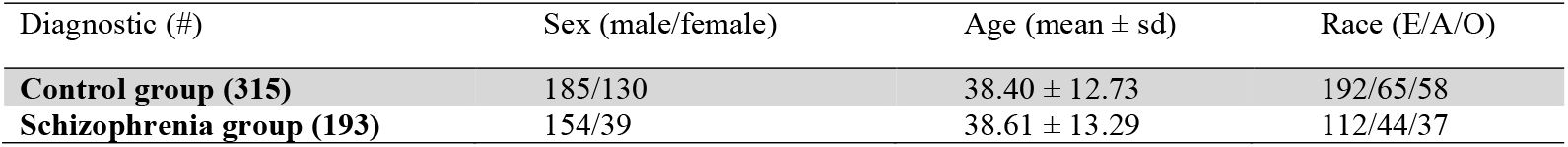
Demographic information. E: European, A: American, O: Other.

| Diagnostic (#) | Sex (male/female) | Age (mean ± sd) | Race (E/A/O) |
| --- | --- | --- | --- |
| **Control group (315)** | 185/130 | 38.40 ± 12.73 | 192/65/58 |
| **Schizophrenia group (193)** | 154/39 | 38.61 ± 13.29 | 112/44/37 |

We observed system-wide disruptions in brain dynamic functional integration (**Fig. 3**). We identified aberrant dynamic spFNC across all the above-mentioned networks in schizophrenia, among which the spFNC pairs of the CER, TEMP, and MTR-S are affected the most. As hypothesized, the dynamic spatial coupling between the pDMN and SN reveals sex-specific changes in schizophrenia. The sex-specific effect also exists in dynamic spFNC pairs of “CER/ MTR-S” and “SUB/FRNT.” The sex-specific effect of schizophrenia was only significant in State 1, the state with the highest level of system-wide functional integration.

**Fig. 3.**
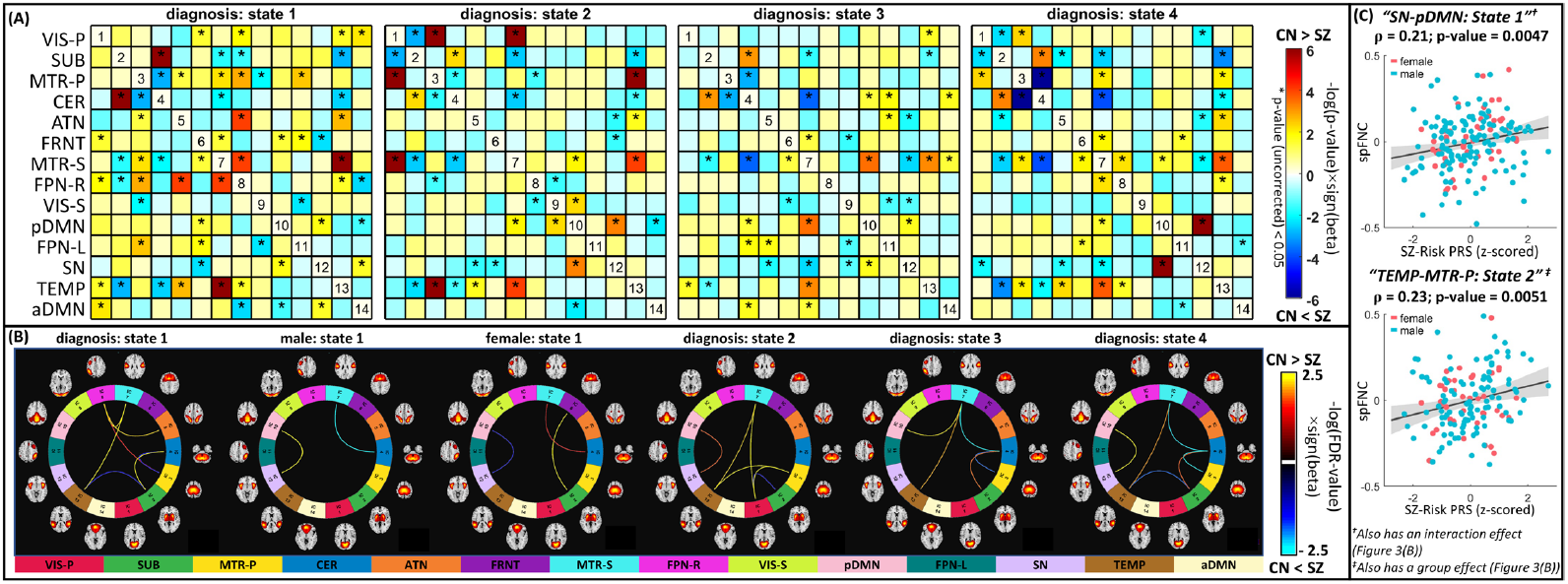
Global Spatial Dynamic Disruption in Schizophrenia (SZ) and Association with Genomic Data. (A) Shows the diagnosis effect (before correcting for multiple comparisons), i.e., SZ vs. control (CN), of the identified spatial functional network connectivity (spFNC) dynamic states. The statistical comparison for each per-state spFNC pair was conducted using a generalized linear regression model (GLM) with age, sex, mean framewise displacement (meanFD), and site as confounding factors and diagnosis and sex-by-diagnosis interactions as predictors of interest. Asterisks (*) represent p-value < 0.05. (B) Connectograms of state spFNC pairs with a significant diagnosis or interaction (diagnosis-by-sex) effects after FDR corrections. Three spFNC pairs from State 1 show significant sex-diagnosis interaction effects, including posterior default mode and salience networks. (C) The association between schizophrenia genetic risk and aberrant dynamic spatial coupling. Among dynamic spFNC pairs with a significant schizophrenia effect, two show significant associations with the polygenic risk score (PRS) after FDR correction. These two include the spFNC between the posterior default mode and salience networks in State 1 with a sex-specific schizophrenia effect and the spFNC between the temporal and primary motor networks in State 2, which show disruption in schizophrenia but with no significant sex effect. SUB: subcortical, VIS-P: visual-primary, VIS-S: visual-secondary, MTR-P: Motor-primary, MTR-S: Motor-secondary, CER: cerebellar, ATN: attention, FRNT: frontal, FPN-R: frontoparietal-right, FPN-L: frontoparietal-left, SN: salience, pDMN: posterior default mode, aDMN: anterior default mode, and TEMP: temporal.

We next evaluated the genomic predisposition of aberrant system-wide dynamic functional integration in schizophrenia. For this purpose, we focused on the schizophrenia-risk single nucleotide polymorphisms (SNPs) residing in the 287 loci reported by a recent large-scale schizophrenia genomic study [40] and computed the polygenic risk score (PRS) for Schizophrenia (SZ) pruned at r-squared < 0.1 [41] using PRSice [42]. The associations between the PRS and aberrant dynamic spFNC were performed on subset of data (European/American = 304/109, schizophrenia/control = 156/257)using Pearson correlation while controlling for diagnosis, sex, age, meanFD, and site and correcting using the FDR. Two dynamic spFNC pairs exhibit significant correlations with the PRS for SZ (PRS-SZ) after FDR corrections, including “pDMN/SN” spFNC in State 1 with a sex-specific schizophrenia effect and “TEMP/MTR-P” the spFNC in State 2 with a significant diagnosis effect (**Fig. 3(C)**). We further investigated these associations separately in European and American populations and observed consistent associations with comparable effect sizes.

### Low Regional Contribution Linked to High Informational Content

Next, we investigated how schizophrenia-related alterations in low-dimensional spatial dynamic states link to diagnosis differences in high-dimensional voxel space, using network spatial maps involved in the two dynamic spFNC pairs with both significant schizophrenia effects and genomic associations. Voxel-wise statistical comparisons are conducted on the networks’ state spatial maps using the same GLM as for spFNC analysis, resulting in one spatial map of beta coefficients for each variable of interest. Subsequently, we computed the spatial similarity between each beta-spatial map and the spatial maps of networks involved in a given spFNC pair using Pearson correlation and compared it with the spatial similarity estimated from null data with the same level of spatial smoothing.

Our results suggest that (1) the aberrations of brain networks reflect changes in dynamic spFNC; (2) schizophrenia affects brain networks in a nuanced and distributed manner across the entire brain; (3) regions with lower network contributions demonstrate a more prominent effect; and (4) the impact of schizophrenia may not necessarily be strongly evident for single voxels, despite significant whole-brain effects.

**Fig. 4(A)** displays the outcomes of the spatial similarity between the beta-spatial maps of variables of interest (e.g., diagnosis) and the network maps. **Fig. 4(B)** exhibits the beta-spatial maps with significant spatial similarity. The beta-spatial map of SN State 1 for the interaction effect (sex-by-diagnosis) and diagnosis effect in males (but not for the group diagnosis effect) show significant spatial similarities with the spatial map of the pDMN (**Fig. 4(A)**). Similarly, the beta-spatial maps of the interaction (sex-by-diagnosis) and diagnosis (male) effects for the State 1 pDMN have significant similarities with the spatial map of the SN (**Fig. 4(A)**). These findings bolster the sex-specific schizophrenia effect observed in “pDMN/SN” spFNC in State 1 (**Fig. 3**).

**Fig. 4.**
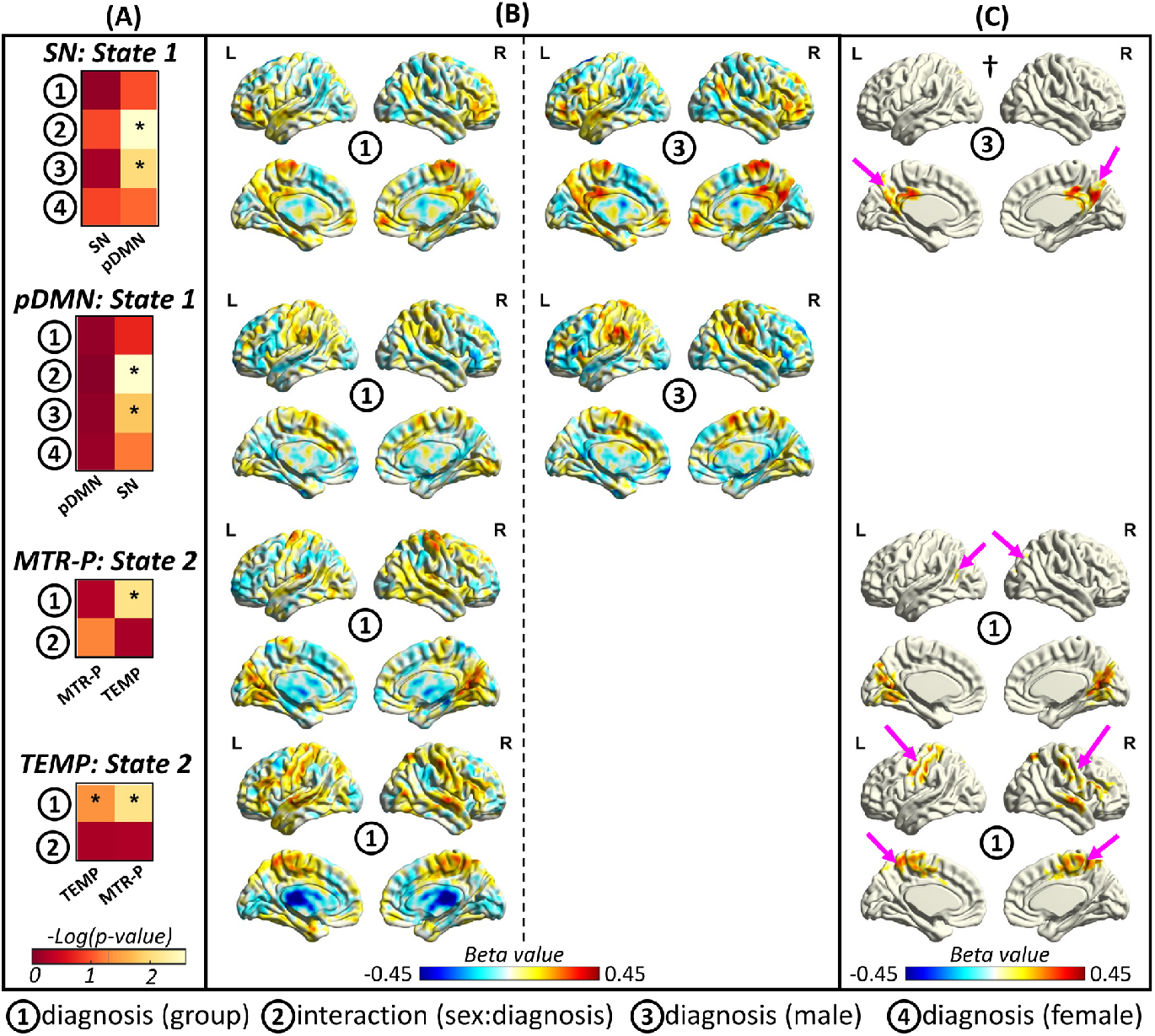
The Evidence of Aberrant Dynamic Spatial Coupling on Network Dynamic Spatial Maps. We conducted voxel-wise statistical analysis on the spatial patterns of the networks to evaluate if the effect of schizophrenia on dynamic spFNC emerges in the network dynamic spatial maps. We focus on the spatial patterns involved in the two spFNC pairs with significant schizophrenia effects and genomic associations. **(A)** shows the results of the spatial similarity (measured by Pearson correlation) between the spatial map of a specific network for a given state and the beta-spatial maps of the variables of interest (e.g., diagnosis) from voxel-wise statistical analysis. Asterisks represent those with significant spatial similarity (p-value < 0.05). The results are aligned with dynamic spFNC findings. For instance, we observe significant spatial similarity with the posterior default mode network (pDMN) for the interaction effect in the salience network (SN) state 1. Another example is the spatial map of the diagnosis group effect (but not the interaction effect) obtain using voxel-wise analysis of the primary somatomotor network (MTR-P) in State 2, which shows significant spatial similarity with the spatial map of the temporal (TEMP) network. **(B)** shows the spatial maps of variables of interest with significant spatial similarity. **(C)** shows the cluster of voxels that survived cluster-wise correction. Pink arrows illustrate clusters in which the paired networks have a strong contribution. For example, the pink arrows in † show the cluster in the posterior cingulate cortex (PCC) and precuneus that survived cluster-wise correction. The PCC and precuneus are the cores of the pDMN and contribute significantly to the pDMN.

For the State 1 pDMN, the impact of schizophrenia was not strong enough to manifest at the voxel level (**Fig. 4(C))**, despite the significant whole-brain spatial similarities between the beta-spatial maps of the interactions and male diagnosis effects. However, the analysis of the SN map in State 1 revealed significant interaction and male diagnosis effects in regions commonly associated with the pDMN (i.e., including the posterior cingulate cortex and precuneus) (**Fig. 4(C)**). It is essential to emphasize that these regions are often masked out in previous analyses due to their low contribution (i.e., weak functional connectivity) to the SN. Therefore, the sex-related changes in schizophrenia that we detected here were previously overlooked in previous studies.

For the MTR-P and TEMP networks in State 2 with significant diagnostic group effects in their spFNC (**Fig. 3**), the beta-spatial map of the diagnosis group effect shows significant spatial similarity with the TEMP and MTR-P networks, respectively (**Fig. 4**(A)). At the voxel level, regions with significant diagnosis group effects for the TEMP network resemble the MTR-P network (**Fig. 4(C))**. For the MTR-P network, the diagnosis group effect spatial map contains a brain area with a significant contribution to TEMP but also contains regions of the VIS-P, which also show aberrant FNC with the MTR-P in State 2.

We also conducted a voxel-wise analysis to evaluate the associations between the PRS and state network spatial maps for two selected dynamic FNC pairs. Similar to earlier genomic association analysis, we used Pearson correlation while controlling for diagnosis, sex, age, meanFD, and site. The genomic association maps reveal significant spatial similarities (p-value < 0.05) with the paired networks for all four evaluated state networks. For example, the genomic association map for the pDMN in State 1 shows significant spatial similarity with the SN map (p-value = 0.03).

## Discussion

Information about dynamic spatial dependence, which has been overlooked, holds immense potential to substantially impact the clinical landscape as it quantifies the continuous integration and segregation of brain networks. Particularly, it can advance our understanding of schizophrenia, a disorder often characterized by dysconnectivity and disruptions in system-wide functional integration.

This study presents a reference-informed network estimation technique to capture time-resolved spatial brain networks and dynamic spatial dependence. This technique not only effectively estimates time-resolved networks from short and low SNR time segments but also controls for the influence of other networks in the estimations. This has a potential advantage over our previous work [9], which utilized sliding window correlation as an alternative approach. Furthermore, while our previous effort [43] focused solely on the high-order statistical portion of spFNC (due to technical limitations), this new method provides a more generalizable and computationally efficient method that is also capable of capturing information about the spatial dependence related to the second order and higher order statistics. By doing so, we can more accurately quantify the dynamic integration and segregation, enhancing our understanding of dysconnectivity in brain dynamics associated with schizophrenia.

We identified four distinct spatial dynamic states with unique network integration and segregation patterns, collectively providing a low-dimensional summary of global brain state dynamics. State 4 emerged as the most segregated state with the highest rate of between-state transitioning. Moreover, the brain spends more time in this state than in others. We propose that this state functions as a hub-like transition state. However, all states have a similar life expectancy, suggesting that State 4 is not a steady state. These findings must be verified in an independent dataset to ensure replicability and generalizability.

We also examined the impact of schizophrenia on low-dimensional spatial dynamic states. Inferring from the dysconnectivity hypothesis [44], disturbances in behavior and psychopathology are expected to be associated with disruptions in the reconfiguration of system-wide brain functional integration. Our findings support this proposition by demonstrating significant alterations in the dynamics of spatial coupling in both men and women with schizophrenia.

In general, individuals with schizophrenia present lower spFNC integration across all states and most networks. Specifically, the patterns of lower integration were more widespread in the MTRs and PFN-R (**Fig. 3)**. This is in line with previous reports of global connectivity deficits in schizophrenia, with more prominent alterations in the frontal and temporal regions [45, 46]. We also found that individuals with schizophrenia present, to a lesser extent, higher spFNC integration in specific pairs of networks, mainly involving the SUB, TEMP, MTRs, CER networks (**Fig. 3**). This is also in line with previous reports of functional hyperconnectivity in individuals with schizophrenia [47-49].

While widespread disruption in networks’ spatial coupling occurs across the whole brain and all dynamic states, the highly integrated, modular State 1 shows a great number of dysconnectivity patterns. The alterations observed in the “pDMN/SN” and “FPN-R/ATN” spFNC in this state may be related to disturbances in cognitive and psychopathology associated with psychosis [25, 50]. These findings are in line with the triple-network (TPN) model [50], suggesting the abnormal striatal dopamine release may lead to disruptions in the dynamics among the DMN, SN, and FPN, thus potentially contributing to the misattribution of salience to irrelevant external stimuli and self-referential mental events.

Moreover, functional connectivity aberrations in the pDMN have been repeatedly involved with schizophrenia pathophysiology [51-53]. Our results support and extend these findings by showing that disruptions in “pDMN/SN” spFNC are present in all four dynamic states (although this did not survive FDR correction in State 3). We also found a sex-by-diagnosis interaction effect for “pDMN/SN” spFNC in State 1, which could explain why some previous work found mixed results [52, 54-56], or no alterations [57] in the DMN functional connectivity of individuals with schizophrenia. This pair also shows an association with schizophrenia PRS, adding another confirmation layer and suggesting a potential sex-specific psychosis neurobiological marker. However, the reproducibility of these results should be verified through an independent dataset.

Our findings suggest some spFNC pairs present statistically significant differences related to schizophrenia that are fleeting, temporally localized to only a few states (i.e., exhibit a state-like property), which contrasts with spFNC pairs where the SZ-changes are consistently present regardless of the brain state (a trait-like property). The PFN-R/ATN coupling exemplifies a state-like spFNC pair, whereas the TEMP/MTR-S pair is a notable trait-like example. Intriguingly, based on Neurosynth (https://www.neurosynth.org/) [58], the highest activation point for the TEMP is linked to semantic integration and language, while the peak activation for MTR-S is associated with speech production. Disruptions in language-related regions and networks in psychotic disorders are a finding well-established by prior studies and might be linked to auditory verbal hallucinations [59-64]. It should be noted that other pairs, such as pDMN/SN (with diagnosis-by-sex interaction effect), MTR-S/SUB, MTR-S/CER, and SUB/CER, demonstrate a similar trait-like pattern, although the difference did not survive multiple comparison corrections in all states (**Fig 3(A)**). Importantly the SZ-changes in MTR-S/SUB/CER support the cerebellar-thalamic-cortical dysconnectivity model of psychosis [65-67].

Another major finding is the spatial decoupling between TEMP and MTR networks across all states, among which TEMP/MTR-P spFNC in State 2 is correlated with PRS-SZ. Alterations in functional connectivity in temporal, somatosensory, and motor regions have been previously associated with schizophrenia [24], but to the best of our knowledge, this is the first study to show an association between those aberrations and PRS-SZ. It is known that individuals with schizophrenia present motor impairments such as delayed motor milestones, poor coordination, or sensory deficiencies (neurologic soft signs) [68]. These soft signs have been suggested to be present since infancy, before illness onset, and associated with a genetic vulnerability that leads to neurodevelopmental disruptions [69]. It has also been proposed that temporal cortex dysfunction plays a role in social cognition and theory of mind impairments in schizophrenia due to difficulties detecting subtle emotional components of auditory inputs, leading to reduced social interaction skills and marked deficiencies in psychosocial functioning [70]. Additionally, the PRS-SZ reflects the overall genetic risk burden of 287 SZ-related loci, for which a fine mapping revealed a diverse set of synaptic proteins and suggested that multiple functional interactions of SZ risk converge on synapses [40]. It was also noted that the 2,196 genes mapped to the 287 risk loci of schizophrenia showed a significant enrichment (Fisher’s exact test, p = 2.61E-09) in 278 human accelerated region-genes (HAR-genes) that had been found to ultimately impact the evolution and development of human high cognitive traits [71]. While our findings suggest a relationship between genetic risk for schizophrenia and TEMP/MTR networks, it is intriguing to investigate whether this genetic predisposition could lead to synaptic alterations in these networks during neurodevelopment, resulting in poor functional integration and the early emergence of neurological signs.

A significant breakthrough has been made regarding the significance of regions with weak functional connectivity (**Fig. 4**). Our findings reveal that the distortion in spatial coupling is embedded in high-dimensional (voxel-level) space in brain regions with low contributions to the corresponding networks (**Fig. 4**). For instance, significant SZ interaction and male diagnosis effects in the SN State 1 are exclusively exhibited in regions with a low contribution to the SN. This observation is significant because current research often overlooks regions with small contributions during voxel-wise analysis (i.e., use a mask of regions with high amplitude in spatial maps). This complements our recent finding [26] on the importance of time points with low contributions in capturing schizophrenia-related changes, calling for further investigation.

## Limitations and Future Direction

This study made choices based on existing knowledge from previous studies [16, 19, 72]. However, the impact of different choices and the generalizability of clinical findings should be assessed by future studies, particularly considering the inconsistency among previous schizophrenia-related findings. From the method perspective, future studies should investigate the impact of window lengths and spatial smoothing [4].

While MOO-ICAR is a good tool for capturing brain networks from small data lengths [4], its spatially constrained nature and imposed independence may limit our ability to fully capture spatial dynamics and subject-specific properties. Therefore, future studies should develop new approaches that are optimized to enable a more precise and comprehensive estimation of time-resolved subject-specific networks. By advancing in this direction, researchers can enhance our understanding of the intricate dynamics of the brain and obtain more accurate insights into individual brain-function variations.

It is essential to recognize that various factors within this sample, such as medication status and symptom severity, may influence the results and limit their applicability to other populations or specific subgroups. Thus, it is imperative to utilize an independent and more homogeneous dataset with larger sample sizes to assess the replicability and generalizability of these findings with a particular focus on medication use and symptomatic manifestations.

While our post hoc analysis found no statistically significant association between medication (chlorpromazine equivalence scores) and two dynamic spFNC pairs with significant schizophrenia effects and genomic associations, it is critical to evaluate the impact of medication on SZ-related changes in brain dynamics.

For example, future studies could investigate the effect of successful/unsuccessful pharmacological treatments on aberrant brain dynamic spatial couplings in temporal and somatosensory networks for individuals with auditory hallucinations. Additionally, future research should evaluate our findings in other cohorts, such as first-episode psychosis, individuals at high risk of psychosis, or first-degree relatives.

While the neurodevelopmental hypothesis of schizophrenia is widely accepted, our study cannot establish causality. However, the correlation between PRS-SZ and two specific aberrations in spFNC integration suggests that genetic vulnerability may influence these alterations. Longitudinal studies with larger datasets are needed to understand the neurodevelopmental trajectory of schizophrenia.

## Acknowledgments

This work was supported by grants from the National Institutes of Health grant numbers R01MH123610, R01EB020407, R01MH118695, and NSF 2112455 to Dr. Vince D. Calhoun and 5R01MH119251 to Dr. Armin Iraji.

## References

[1] A. Iraji, R. Miller, T. Adali, and V. D. Calhoun, “Space: A Missing Piece of the Dynamic Puzzle,” (in eng), Trends Cogn Sci, vol. 24, no. 2, pp. 135–149, Feb 2020, doi: 10.1016/j.tics.2019.12.004.

[2] E. A. Allen et al., “A baseline for the multivariate comparison of resting-state networks,” (in eng), Front Syst Neurosci, vol. 5, p. 2, 2011, doi: 10.3389/fnsys.2011.00002.

[3] Y. Du et al., “NeuroMark: An automated and adaptive ICA based pipeline to identify reproducible fMRI markers of brain disorders,” (in eng), Neuroimage Clin, vol. 28, p. 102375, 2020, doi: 10.1016/j.nicl.2020.102375.

[4] A. Iraji et al., “Canonical and Replicable Multi-Scale Intrinsic Connectivity Networks in 100k+ Resting-State fMRI Datasets,” bioRxiv, p. 2022.09.03.506487, 2022, doi: 10.1101/2022.09.03.506487.

[5] A. Schaefer et al., “Local-Global Parcellation of the Human Cerebral Cortex from Intrinsic Functional Connectivity MRI,” (in eng), Cereb Cortex, vol. 28, no. 9, pp. 3095–3114, Sep 1 2018, doi: 10.1093/cercor/bhx179.

[6] R. C. Craddock, G. A. James, P. E. Holtzheimer, 3rd, X. P. Hu, and H. S. Mayberg, “A whole brain fMRI atlas generated via spatially constrained spectral clustering,” (in eng), Hum Brain Mapp, vol. 33, no. 8, pp. 1914–28, Aug 2012, doi: 10.1002/hbm.21333.

[7] B. T. Yeo et al., “The organization of the human cerebral cortex estimated by intrinsic functional connectivity,” (in eng), J Neurophysiol, vol. 106, no. 3, pp. 1125–65, Sep 2011, doi: 10.1152/jn.00338.2011.

[8] V. D. Calhoun, R. Miller, G. Pearlson, and T. Adali, “The chronnectome: time-varying connectivity networks as the next frontier in fMRI data discovery,” (in eng), Neuron, vol. 84, no. 2, pp. 262–74, Oct 22 2014, doi: 10.1016/j.neuron.2014.10.015.

[9] A. Iraji et al., “The spatial chronnectome reveals a dynamic interplay between functional segregation and integration,” (in eng), Hum Brain Mapp, vol. 40, no. 10, pp. 3058–3077, Jul 2019, doi: 10.1002/hbm.24580.

[10] A. Iraji et al., “Spatial dynamics within and between brain functional domains: A hierarchical approach to study time-varying brain function,” (in eng), Hum Brain Mapp, vol. 40, no. 6, pp. 1969–1986, Apr 15 2019, doi: 10.1002/hbm.24505.

[11] S. Bhinge, Q. Long, V. D. Calhoun, and T. Adali, “Spatial Dynamic Functional Connectivity Analysis Identifies Distinctive Biomarkers in Schizophrenia,” (in eng), Front Neurosci, vol. 13, p. 1006, 2019, doi: 10.3389/fnins.2019.01006.

[12] A. Boukhdhir, Y. Zhang, M. Mignotte, and P. Bellec, “Unraveling reproducible dynamic states of individual brain functional parcellation,” (in eng), Netw Neurosci, vol. 5, no. 1, pp. 28–55, 2021, doi: 10.1162/netn_a_00168.

[13] L. Q. Uddin et al., “Controversies and current progress on large-scale brain network nomenclature from OHBM WHATNET: Workgroup for HArmonized Taxonomy of NETworks,” 2022.

[14] V. D. Calhoun, T. Adali, G. D. Pearlson, and J. J. Pekar, “A method for making group inferences from functional MRI data using independent component analysis,” (in eng), Hum Brain Mapp, vol. 14, no. 3, pp. 140–51, Nov 2001, doi: 10.1002/hbm.1048.

[15] A. Iraji et al., “The connectivity domain: Analyzing resting state fMRI data using feature-based data-driven and model-based methods,” (in eng), Neuroimage, vol. 134, pp. 494–507, Jul 1 2016, doi: 10.1016/j.neuroimage.2016.04.006.

[16] A. Iraji, A. Faghiri, N. Lewis, Z. Fu, S. Rachakonda, and V. D. Calhoun, “Tools of the trade: estimating time-varying connectivity patterns from fMRI data,” (in eng), Soc Cogn Affect Neurosci, vol. 16, no. 8, pp. 849–874, Aug 5 2021, doi: 10.1093/scan/nsaa114.

[17] Y. Du and Y. Fan, “Group information guided ICA for fMRI data analysis,” (in eng), Neuroimage, vol. 69, pp. 157–97, Apr 1 2013, doi: 10.1016/j.neuroimage.2012.11.008.

[18] Y. Du, E. A. Allen, H. He, J. Sui, L. Wu, and V. D. Calhoun, “Artifact removal in the context of group ICA: A comparison of single-subject and group approaches,” (in eng), Hum Brain Mapp, vol. 37, no. 3, pp. 1005–25, Mar 2016, doi: 10.1002/hbm.23086.

[19] E. A. Allen, E. Damaraju, S. M. Plis, E. B. Erhardt, T. Eichele, and V. D. Calhoun, “Tracking whole-brain connectivity dynamics in the resting state,” (in eng), Cereb Cortex, vol. 24, no. 3, pp. 663–76, Mar 2014, doi: 10.1093/cercor/bhs352.

[20] A. Iraji et al., “Multi-spatial-scale dynamic interactions between functional sources reveal sex-specific changes in schizophrenia,” Network Neuroscience, vol. 6, no. 2, pp. 357–381, 2022, doi: 10.1162/netn_a_00196.

[21] J. W. Buckholtz and A. Meyer-Lindenberg, “Psychopathology and the human connectome: toward a transdiagnostic model of risk for mental illness,” (in eng), Neuron, vol. 74, no. 6, pp. 990–1004, Jun 21 2012, doi: 10.1016/j.neuron.2012.06.002.

[22] K. J. Friston, “Dysfunctional connectivity in schizophrenia,” (in eng), World Psychiatry, vol. 1, no. 2, pp. 66–71, Jun 2002.

[23] V. D. Calhoun, J. Sui, K. Kiehl, J. Turner, E. Allen, and G. Pearlson, “Exploring the psychosis functional connectome: aberrant intrinsic networks in schizophrenia and bipolar disorder,” (in eng), Front Psychiatry, vol. 2, p. 75, 2011, doi: 10.3389/fpsyt.2011.00075.

[24] E. Damaraju et al., “Dynamic functional connectivity analysis reveals transient states of dysconnectivity in schizophrenia,” (in eng), Neuroimage Clin, vol. 5, pp. 298–308, 2014, doi: 10.1016/j.nicl.2014.07.003.

[25] S. M. Hare et al., “Salience-Default Mode Functional Network Connectivity Linked to Positive and Negative Symptoms of Schizophrenia,” (in eng), Schizophr Bull, vol. 45, no. 4, pp. 892–901, Jun 18 2019, doi: 10.1093/schbul/sby112.

[26] A. Iraji et al., “Moving beyond the ‘CAP’ of the Iceberg: Intrinsic connectivity networks in fMRI are continuously engaging and overlapping,” (in eng), Neuroimage, vol. 251, p. 119013, May 1 2022, doi: 10.1016/j.neuroimage.2022.119013.

[27] S. Li et al., “Dysconnectivity of Multiple Brain Networks in Schizophrenia: A Meta-Analysis of Resting-State Functional Connectivity,” (in eng), Front Psychiatry, vol. 10, p. 482, 2019, doi: 10.3389/fpsyt.2019.00482.

[28] D. Dong, Y. Wang, X. Chang, C. Luo, and D. Yao, “Dysfunction of Large-Scale Brain Networks in Schizophrenia: A Meta-analysis of Resting-State Functional Connectivity,” (in eng), Schizophr Bull, vol. 44, no. 1, pp. 168–181, Jan 13 2018, doi: 10.1093/schbul/sbx034.

[29] Q. Long, S. Bhinge, V. D. Calhoun, and T. Adali, “Graph-theoretical analysis identifies transient spatial states of resting-state dynamic functional network connectivity and reveals dysconnectivity in schizophrenia,” (in eng), J Neurosci Methods, vol. 350, p. 109039, Feb 15 2021, doi: 10.1016/j.jneumeth.2020.109039.

[30] Q. Long, S. Bhinge, V. D. Calhoun, and T. Adali, “Relationship between Dynamic Blood-Oxygen-Level-Dependent Activity and Functional Network Connectivity: Characterization of Schizophrenia Subgroups,” (in eng), Brain Connect, vol. 11, no. 6, pp. 430–446, Aug 2021, doi: 10.1089/brain.2020.0815.

[31] V. Pugliese et al., “Aberrant salience correlates with psychotic dimensions in outpatients with schizophrenia spectrum disorders,” (in eng), Ann Gen Psychiatry, vol. 21, no. 1, p. 25, Jul 3 2022, doi: 10.1186/s12991-022-00402-5.

[32] R. Li, X. Ma, G. Wang, J. Yang, and C. Wang, “Why sex differences in schizophrenia?,” (in eng), J Transl Neurosci (Beijing), vol. 1, no. 1, pp. 37–42, Sep 2016.

[33] A. Riecher-Rössler, S. Butler, and J. Kulkarni, “Sex and gender differences in schizophrenic psychoses-a critical review,” (in eng), Arch Womens Ment Health, vol. 21, no. 6, pp. 627–648, Dec 2018, doi: 10.1007/s00737-018-0847-9.

[34] B. Carter, J. Wootten, S. Archie, A. L. Terry, and K. K. Anderson, “Sex and gender differences in symptoms of early psychosis: a systematic review and meta-analysis,” (in eng), Arch Womens Ment Health, vol. 25, no. 4, pp. 679–691, Aug 2022, doi: 10.1007/s00737-022-01247-3.

[35] A. Mendrek and A. Mancini-MarÏe, “Sex/gender differences in the brain and cognition in schizophrenia,” (in eng), Neurosci Biobehav Rev, vol. 67, pp. 57–78, Aug 2016, doi: 10.1016/j.neubiorev.2015.10.013.

[36] H. Long et al., “Sex-related Difference in Mental Rotation Performance is Mediated by the special Functional Connectivity Between the Default Mode and Salience Networks,” (in eng), Neuroscience, vol. 478, pp. 65–74, Dec 1 2021, doi: 10.1016/j.neuroscience.2021.10.009.

[37] K. E. Lawrence et al., “Sex Differences in Functional Connectivity of the Salience, Default Mode, and Central Executive Networks in Youth with ASD,” (in eng), Cereb Cortex, vol. 30, no. 9, pp. 5107–5120, Jul 30 2020, doi: 10.1093/cercor/bhaa105.

[38] R. Canitano and M. Pallagrosi, “Autism Spectrum Disorders and Schizophrenia Spectrum Disorders: Excitation/Inhibition Imbalance and Developmental Trajectories,” (in eng), Front Psychiatry, vol. 8, p. 69, 2017, doi: 10.3389/fpsyt.2017.00069.

[39] Y. Benjamini and Y. Hochberg, “Controlling the False Discovery Rate: A Practical and Powerful Approach to Multiple Testing,” Journal of the Royal Statistical Society. Series B (Methodological), vol. 57, no. 1, pp. 289–300, 1995. [Online]. Available: http://www.jstor.org/stable/2346101.

[40] V. Trubetskoy et al., “Mapping genomic loci implicates genes and synaptic biology in schizophrenia,” Nature, vol. 604, no. 7906, pp. 502–508, Apr 2022, doi: 10.1038/s41586-022-04434-5.

[41] V. Trubetskoy et al., “Mapping genomic loci implicates genes and synaptic biology in schizophrenia,” (in eng), Nature, vol. 604, no. 7906, pp. 502–508, Apr 2022, doi: 10.1038/s41586-022-04434-5.

[42] S. W. Choi, T. S. H. Mak, and P. F. O’Reilly, “Tutorial: a guide to performing polygenic risk score analyses,” (in English), Nat Protoc, vol. 15, no. 9, pp. 2759–2772, Sep 2020, doi: 10.1038/s41596-020-0353-1.

[43] S. Ma, V. D. Calhoun, R. Phlypo, and T. Adalı, “Dynamic changes of spatial functional network connectivity in healthy individuals and schizophrenia patients using independent vector analysis,” (in eng), Neuroimage, vol. 90, pp. 196–206, Apr 15 2014, doi: 10.1016/j.neuroimage.2013.12.063.

[44] K. Friston, H. R. Brown, J. Siemerkus, and K. E. Stephan, “The dysconnection hypothesis (2016),” (in eng), Schizophr Res, vol. 176, no. 2-3, pp. 83–94, Oct 2016, doi: 10.1016/j.schres.2016.07.014.

[45] A. Fornito, A. Zalesky, C. Pantelis, and E. T. Bullmore, “Schizophrenia, neuroimaging and connectomics,” (in eng), Neuroimage, vol. 62, no. 4, pp. 2296–314, Oct 1 2012, doi: 10.1016/j.neuroimage.2011.12.090.

[46] M. P. van den Heuvel and A. Fornito, “Brain networks in schizophrenia,” (in eng), Neuropsychol Rev, vol. 24, no. 1, pp. 32–48, Mar 2014, doi: 10.1007/s11065-014-9248-7.

[47] H. Liu et al., “Schizophrenic patients and their unaffected siblings share increased resting-state connectivity in the task-negative network but not its anticorrelated task-positive network,” (in eng), Schizophr Bull, vol. 38, no. 2, pp. 285–94, Mar 2012, doi: 10.1093/schbul/sbq074.

[48] S. Walther, K. Stegmayer, A. Federspiel, S. Bohlhalter, R. Wiest, and P. V. Viher, “Aberrant Hyperconnectivity in the Motor System at Rest Is Linked to Motor Abnormalities in Schizophrenia Spectrum Disorders,” (in eng), Schizophr Bull, vol. 43, no. 5, pp. 982–992, Sep 1 2017, doi: 10.1093/schbul/sbx091.

[49] M. Avram, F. Brandl, J. Bäuml, and C. Sorg, “Cortico-thalamic hypo- and hyperconnectivity extend consistently to basal ganglia in schizophrenia,” (in eng), Neuropsychopharmacology, vol. 43, no. 11, pp. 2239–2248, Oct 2018, doi: 10.1038/s41386-018-0059-z.

[50] V. Menon, L. Palaniyappan, and K. Supekar, “Integrative Brain Network and Salience Models of Psychopathology and Cognitive Dysfunction in Schizophrenia,” (in eng), Biol Psychiatry, Oct 4 2022, doi: 10.1016/j.biopsych.2022.09.029.

[51] M. L. Hu et al., “A Review of the Functional and Anatomical Default Mode Network in Schizophrenia,” (in eng), Neurosci Bull, vol. 33, no. 1, pp. 73–84, Feb 2017, doi: 10.1007/s12264-016-0090-1.

[52] J. M. Sheffield and D. M. Barch, “Cognition and resting-state functional connectivity in schizophrenia,” (in eng), Neurosci Biobehav Rev, vol. 61, pp. 108–20, Feb 2016, doi: 10.1016/j.neubiorev.2015.12.007.

[53] S. A. Meda et al., “Multivariate analysis reveals genetic associations of the resting default mode network in psychotic bipolar disorder and schizophrenia,” (in eng), Proc Natl Acad Sci U S A, vol. 111, no. 19, pp. E2066–75, May 13 2014, doi: 10.1073/pnas.1313093111.

[54] P. Skudlarski et al., “Brain connectivity is not only lower but different in schizophrenia: a combined anatomical and functional approach,” (in eng), Biol Psychiatry, vol. 68, no. 1, pp. 61–9, Jul 1 2010, doi: 10.1016/j.biopsych.2010.03.035.

[55] G. Mingoia et al., “Default mode network activity in schizophrenia studied at resting state using probabilistic ICA,” (in eng), Schizophr Res, vol. 138, no. 2–3, pp. 143–9, Jul 2012, doi: 10.1016/j.schres.2012.01.036.

[56] F. Orliac et al., “Links among resting-state default-mode network, salience network, and symptomatology in schizophrenia,” (in eng), Schizophr Res, vol. 148, no. 1-3, pp. 74–80, Aug 2013, doi: 10.1016/j.schres.2013.05.007.

[57] N. D. Wolf et al., “Dysconnectivity of multiple resting-state networks in patients with schizophrenia who have persistent auditory verbal hallucinations,” (in eng), J Psychiatry Neurosci, vol. 36, no. 6, pp. 366–74, Nov 2011, doi: 10.1503/jpn.110008.

[58] T. Yarkoni, R. A. Poldrack, T. E. Nichols, D. C. Van Essen, and T. D. Wager, “Large-scale automated synthesis of human functional neuroimaging data,” (in eng), Nat Methods, vol. 8, no. 8, pp. 665–70, Jun 26 2011, doi: 10.1038/nmeth.1635.

[59] L. E. DeLisi, “Speech disorder in schizophrenia: review of the literature and exploration of its relation to the uniquely human capacity for language,” (in eng), Schizophr Bull, vol. 27, no. 3, pp. 481–96, 2001, doi: 10.1093/oxfordjournals.schbul.a006889.

[60] X. Chang et al., “Resting-state functional connectivity in medication-naÏve schizophrenia patients with and without auditory verbal hallucinations: A preliminary report,” (in eng), Schizophr Res, vol. 188, pp. 75–81, Oct 2017, doi: 10.1016/j.schres.2017.01.024.

[61] L. B. Cui et al., “Putamen-related regional and network functional deficits in first-episode schizophrenia with auditory verbal hallucinations,” (in eng), Schizophr Res, vol. 173, no. 1-2, pp. 13–22, May 2016, doi: 10.1016/j.schres.2016.02.039.

[62] K. T. Mueser, A. S. Bellack, and E. U. Brady, “Hallucinations in schizophrenia,” (in eng), Acta Psychiatr Scand, vol. 82, no. 1, pp. 26–9, Jul 1990, doi: 10.1111/j.1600-0447.1990.tb01350.x.

[63] S. McCarthy-Jones et al., “Occurrence and co-occurrence of hallucinations by modality in schizophrenia-spectrum disorders,” (in eng), Psychiatry Res, vol. 252, pp. 154–160, Jun 2017, doi: 10.1016/j.psychres.2017.01.102.

[64] D. F. Salisbury, J. Kohler, M. E. Shenton, and R. W. McCarley, “Deficit Effect Sizes and Correlations of Auditory Event-Related Potentials at First Hospitalization in the Schizophrenia Spectrum,” (in eng), Clin EEG Neurosci, vol. 51, no. 4, pp. 198–206, Jul 2020, doi: 10.1177/1550059419868115.

[65] N. D. Woodward, H. Karbasforoushan, and S. Heckers, “Thalamocortical dysconnectivity in schizophrenia,” (in eng), Am J Psychiatry, vol. 169, no. 10, pp. 1092–9, Oct 2012, doi: 10.1176/appi.ajp.2012.12010056.

[66] Y. Wei et al., “Aberrant Cerebello-Thalamo-Cortical Functional and Effective Connectivity in First-Episode Schizophrenia With Auditory Verbal Hallucinations,” (in eng), Schizophr Bull, vol. 48, no. 6, pp. 1336–1343, Nov 18 2022, doi: 10.1093/schbul/sbab142.

[67] H. Cao, M. Ingvar, C. M. Hultman, and T. Cannon, “Evidence for cerebello-thalamo-cortical hyperconnectivity as a heritable trait for schizophrenia,” (in eng), Transl Psychiatry, vol. 9, no. 1, p. 192, Aug 20 2019, doi: 10.1038/s41398-019-0531-5.

[68] R. C. Chan, T. Xu, R. W. Heinrichs, Y. Yu, and Y. Wang, “Neurological soft signs in schizophrenia: a meta-analysis,” (in eng), Schizophr Bull, vol. 36, no. 6, pp. 1089–104, Nov 2010, doi: 10.1093/schbul/sbp011.

[69] I. Bombin, C. Arango, and R. W. Buchanan, “Significance and meaning of neurological signs in schizophrenia: two decades later,” (in eng), Schizophr Bull, vol. 31, no. 4, pp. 962–77, Oct 2005, doi: 10.1093/schbul/sbi028.

[70] D. C. Javitt and R. A. Sweet, “Auditory dysfunction in schizophrenia: integrating clinical and basic features,” (in eng), Nat Rev Neurosci, vol. 16, no. 9, pp. 535–50, Sep 2015, doi: 10.1038/nrn4002.

[71] Y. Wei et al., “Genetic mapping and evolutionary analysis of human-expanded cognitive networks,” Nat Commun, vol. 10, no. 1, p. 4839, 2019.

[72] M. G. Preti, T. A. Bolton, and D. Van De Ville, “The dynamic functional connectome: State-of-the-art and perspectives,” (in eng), NeuroImage, vol. 160, pp. 41–54, Oct 15 2017, doi: 10.1016/j.neuroimage.2016.12.061.

